# Thermal regimes, but not mean temperatures, drive patterns of rapid climate adaptation at a continent-scale: evidence from the introduced European earwig across North America

**DOI:** 10.1101/550319

**Authors:** Jean-Claude Tourneur, Joël Meunier

## Abstract

The recent development of human societies has led to major, rapid and often inexorable changes in the environment of most animal species. Over the last decades, a growing number of studies formulated predictions on the modalities of animal adaptation to climate change, questioning how and at what speed animals should adapt to such changes, discussing the levels of risks imposed by changes in the mean and/or variance of temperatures on animal performance, and exploring the underlying roles of phenotypic plasticity and genetic inheritance. These fundamental predictions, however, remain poorly tested using field data. Here, we tested these predictions using a unique continental-scale data set in the European earwig *Forficula auricularia* L, a univoltine insect introduced in North America one century ago. We conducted a common garden experiment, in which we measured 13 life-history traits in 4158 field-sampled earwigs originating from 19 populations across North America. Our results first demonstrate that in less than 100 generations, this species modified 10 of the 13 measured life-history traits in response to the encountered thermal regimes, defined as a variation of temperatures between seasons or months (here winter-summer and autumn-spring temperatures). We found, however, no response to the overall mean monthly temperatures of the invaded locations. Furthermore, our use of a common garden setup reveals that the observed changes in earwigs’ life-history traits are not mere plastic responses to their current environment, but are either due to their genetic background and/or to the environmental conditions they experienced during early life development. Overall, these findings provide continent-scale support to the claims that adaptation to thermal changes occurs quickly, even in insects with long life cycles, and emphasize the importance of thermal regimes over mean population temperatures in climate adaptation.

## Introduction

The dramatic acceleration of climate change observed over the last decades challenges the ability of resident organisms to track these changes and adapt their life histories accordingly [1–3]. Over the last decades, modelling and theoretical approaches have been developed to better understand the nature and extent of animals’ response to such a climate change [2]. These studies formulated key predictions on how and at what speed animals should adapt to such changes, on the respective importance of an increase in the overall mean temperature and/or seasonality of a population on animal performance, as well as on the underlying roles of phenotypic plasticity and genetic inheritance in adaption [4–13]. For instance, these studies suggest that a rapid adaptation to climate change should be facilitated in organisms with fast development and short life-cycles, as found in many arthropods, whereas it should be more difficult in organisms exhibiting slow development and long life-cycle, as found in many vertebrates. Species should also be less sensitive to changes in seasonality compared to changes in overall mean temperatures when they are endotherms and/or when their entire life-cycle occur within a single season, whereas the opposite pattern is expected when they are ectotherms and/or have a life-cycle encompassing several seasons. Finally, phenotypic plasticity is often considered a keystone of rapid adaptation to environmental changes, whereas fixed and inherited patterns of adaptation are often thought to secondarily derive from the canalization of ancestral plastic variation [14].

Although central in our current understanding of animal’s responses to climate change, these fundamental predictions remain poorly tested in the field [15,16]. This is probably because such field data are difficult to collect, as it typically requires measuring variation in life-history traits across multiple natural populations, over several years, and under different kind of climates. However, a powerful alternative consists in using field data of introduced species that quickly invaded large geographic areas exhibiting a broad diversity of thermal constraints [17,18]. In this study, we present and analyze such a unique field data set in one of these species, the European earwig *Forficula auricularia* Linnaeus (Dermaptera: Forficulidae), after its introduction in North America. This insect exhibits a broad native range extending across Europe, Asia and northern Africa [19] from which it has been introduced to Australia, New Zealand, East Africa, East Indies, Tasmania and North America [20–23]. Its presence in North America was first reported on the Pacific coast in Seattle (WA) in 1907 [24], and then on the Atlantic coast in Newport (RI) in 1911 [25] and in Vancouver (BC) in 1919 [26]. From these introductory foci, *F. auricularia* first spread along the coasts to cover areas ranging from British Columbia to California and from Newfoundland to South Carolina, and then reached the interior of the continent in both United States of America [27] and Canada [28–30]. Given that this species produces only one generation per year [31,32], these historical records reveal that its successful colonization of North America and thus its adaptation to a broad diversity of thermal environments occurred in less than 100 generations.

Because the univoltine life cycle of the European earwig lasts up to 2 years and encompasses all seasons and temperatures [31,33], it has long been thought that annual mean temperatures and/or temperature seasonality could be major constraints in the success of *F. auricularia* invasions [21,34]. However, it remains unclear whether this species can mitigate these thermal constraints, and whether it does so by adapting its life cycle and life-history traits [35,36]. The life cycle of the European earwig generally starts with the emergence of new adults in late spring to early July, with variation among populations. These adults form groups of up to several hundred individuals, in which both males and females typically mate with several partners [28,37,38]. Females then burrow in the ground from mid fall to early winter and build a nest where they lay their first clutch of eggs. After egg laying, females stop their foraging activity and provide extensive forms of egg care until hatching [39–45]. The eggs of this first clutch hatch in spring and mothers remain with their newly hatched larvae for several weeks, during which mothers provide larvae with multiple forms of care [44,46,47] and larvae exhibit forms of sibling cooperation [46,48–50]. A few weeks later, the family unit is naturally disrupted. While larvae continue their development to adults in new social groups, some females produce a second clutch of eggs (i.e. iteroparous as compared to semelparous females), which will also receive pre- and post-hatching care and will hatch in late spring [19,31,36]. All females generally die during the following summer [51].

In this study, we used a common garden experiment to explore how *F. auricularia* responded to the different thermal environments encountered during their North American invasion over the last century, i.e. in less than 100 generations. In particular, we 1) tested whether and how individuals altered their life history traits in response to the thermal constraints of the invaded locations, 2) identified the thermal constraints to which they adapted and 3) investigated the role of phenotypic plasticity in this adaptation. From 1988 to 1995, we field-sampled individuals originating from 19 populations located from the East to the West coasts, maintained them under standard laboratory conditions and measured the properties of the 1^st^ and 2^nd^ clutches produced by each female in terms of egg laying date, egg number, egg development time and number of newly hatched larvae. We also recorded the reproductive strategy of the females (iteroparity versus semelparity), their reproductive outcome (total number of eggs and larvae produced over lifetime), as well as the experimental survival duration of the field-sampled males and females. To identify which thermal constraints the tested earwigs adapted to, we tested whether our measurements could be explained by the results of a principal component analyses (PCA) of the mean monthly temperatures of each population. This process characterizes patterns of variation among populations’ temperatures without *a priori* definitions of their associations, i.e. without predetermining the focus on overall mean temperatures and/or specific thermal regimes (defined as variation of temperatures between seasons or months). If *F. auricularia* individuals adapted their life-cycle and life-history traits to the mean temperatures and/or thermal regimes of the population in which they have been sampled (and if this adaptation is determined by their genetic background and/or early life experience), we predict these traits to covary with the overall mean temperatures and/or variation in seasonal temperatures of their population (i.e. all sampled populations should show different performance in the common garden). Conversely, if earwig life-history traits are independent of the thermal environment of the population in which they have been sampled (i.e. no adaptation) and/or are plastic to their current thermal environment, we predict no apparent association between the traits measured in our field-sampled individuals and the thermal regimes of their populations (i.e. all sampled populations should show similar performance in the common garden).

## Material and methods

### Earwig sampling and laboratory rearing

All *F. auricularia* individuals were collected over 7 years among 19 natural populations located across North America (Figure 1, Table 1). These individuals were mostly collected as adults using wooden traps [35] between July and August, and were immediately setup in glass containers (Mason Jars Company, Erie, Pennsylvania, United States of America) in groups of 20 to 30 individuals. These containers received two sheets of creased toilet paper as resting places for earwigs, and were then transported to the laboratory in Montreal, Canada. Upon their arrival, containers were deposited in a shelf covered by a shelter and maintained under the natural outdoor conditions of Montreal. During their transport and outdoor maintenance, containers received an *ad libitum* amount of carrots and pollen as a food source for earwigs, and were supplied with water by means of a cotton pad regularly soaked in water. This setup allowed earwigs to perform non-controlled mating and to live in groups – just like they do under natural conditions [37,38,43,52].

**Figure 1.**
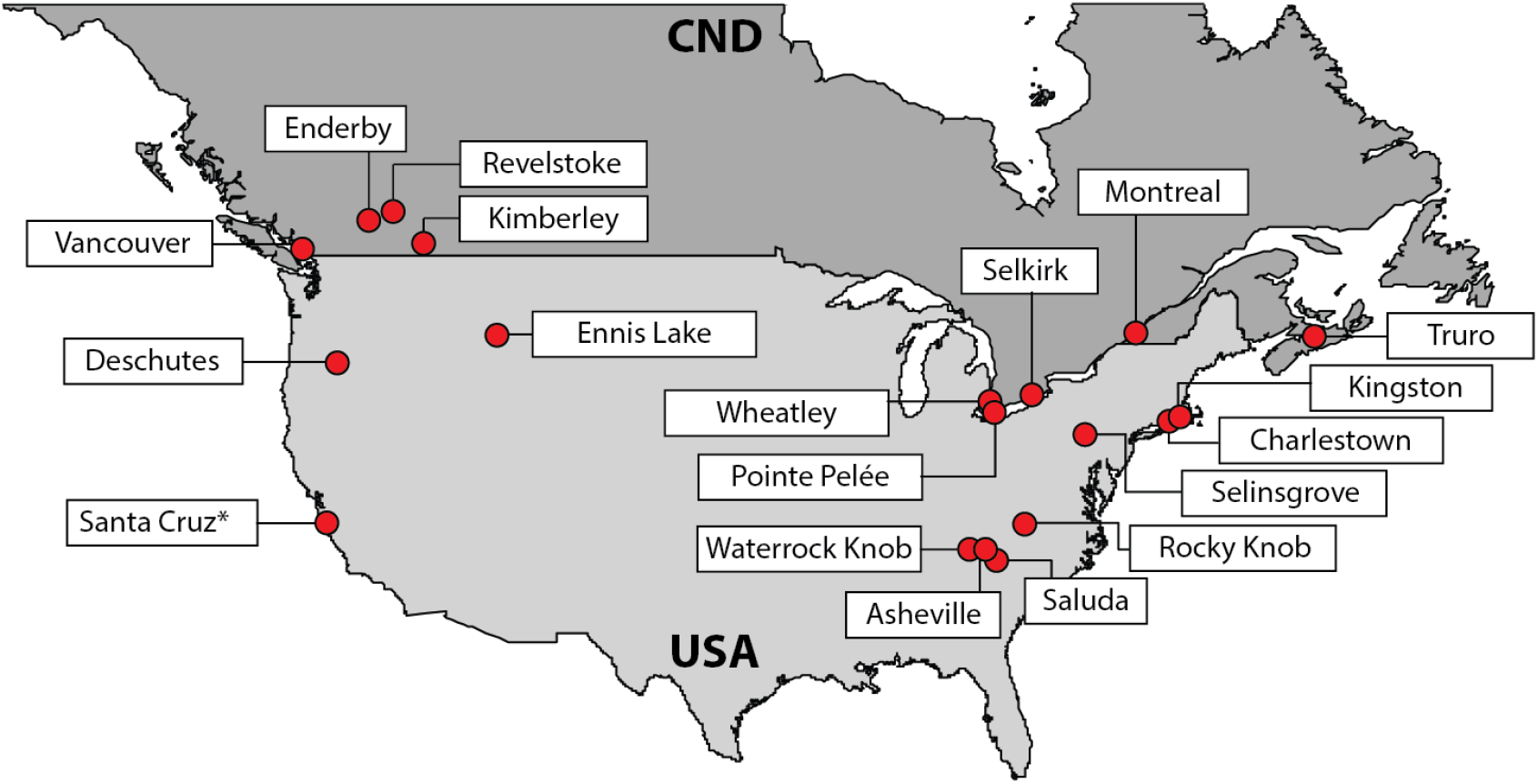
Map showing the 19 sampled populations across Canada (CND) and United States of America (USA). * This population was called San Francisco in Tourneur (2018).

**Table 1.** Details of the 19 sampled populations. The table shows the name and location of each population, their GPS coordinates (Latitude, Longitude), samplings years, total number of mating pair setup across years (N. pairs), and thermal regimes (defined as PC1, PC2 and PC3). * This population was called San Francisco in [35].

| Populations | Country | State (USA)/Province (CDN) | Latitude | Longitude | Samplings | N. pairs | PC1 | PC2 | PC3 |
| --- | --- | --- | --- | --- | --- | --- | --- | --- | --- |
| Asheville | USA | North Carolina | 35.612 | -82.566 | 1994-95 | 80 | 4.60 | 0.73 | 0.74 |
| Charlestown | USA | Rhode Island | 41.383 | -71.642 | 1990 | 42 | 1.37 | 0.73 | -0.84 |
| Deschutes | USA | Oregon | 44.157 | -121.256 | 1990 | 17 | -1.54 | -2.32 | -0.27 |
| Enderby | CDN | British Columbia | 50.551 | -119.14 | 1989-90 | 121 | -1.68 | -0.02 | 1.14 |
| Ennis lake | USA | Montana | 45.447 | -111.695 | 1990 | 36 | -2.86 | -0.36 | 0.02 |
| Kimberley | CDN | British Columbia | 49.635 | -115.998 | 1990 | 94 | -5.27 | -0.95 | 0.73 |
| Kingston | USA | Rhode Island | 41.486 | -71.531 | 1991 | 137 | 1.00 | 0.76 | -0.75 |
| Montreal | CDN | Quebec | 45.542 | -73.893 | 1988,1990-95 | 356 | -2.78 | 2.32 | -0.07 |
| Pointe Pelée | CDN | Ontario | 41.963 | -82.518 | 1992 | 47 | 1.25 | 2.31 | -0.37 |
| Revelstoke | CDN | British Columbia | 50.998 | -118.196 | 1989-90 | 100 | -2.69 | -0.11 | 0.95 |
| Rocky knob | USA | Virginia | 36.832 | -80.345 | 1993-94 | 304 | 1.44 | -0.07 | 0.48 |
| Saluda | USA | North Carolina | 35.198 | -82.353 | 1993-95 | 117 | 5.03 | 0.69 | 0.77 |
| Santa Cruz* | USA | California | 36.926 | -121.845 | 1991 | 130 | 5.04 | -4.50 | -0.57 |
| Selkirk | CDN | Ontario | 42.834 | -79.932 | 1992-94 | 233 | -0.69 | 1.35 | -0.67 |
| Selinsgrove | USA | Pennsylvania | 40.832 | -76.872 | 1993-94 | 134 | 1.76 | 1.83 | 0.27 |
| Truro | CDN | Nova Scotia | 45.372 | -63.264 | 1988 | 39 | -3.46 | -0.13 | -1.42 |
| Vancouver | CDN | British Columbia | 49.252 | -123.24 | 1989,1991 | 84 | 0.23 | -2.89 | -0.05 |
| Waterrock knob | USA | North Carolina | 35.464 | -83.138 | 1991-94 | 167 | -1.88 | -1.69 | 0.22 |
| Wheatley | CDN | Ontario | 42.094 | -82.445 | 1992 | 52 | 1.13 | 2.31 | -0.31 |

One to two months later (between the 7^th^ and the 19^th^ day of October of each year), we used 4158 of these field-sampled individuals to set up 2079 mating pairs (from 17 to 356 pairs per population, see Table 1), in which we subsequently measured 13 life-history traits (see below). These pairs were set up in Petri dishes (diameter 10 cm) lined with a thin layer of moist sand, and in which food was changed and substrate humidified once a week. Each Petri dish was then transferred in a climate chamber and then maintained at 10 ± 1 °C, a temperature close to the overall median temperature of the 19 sampled populations (i.e. 9.5°C, see Table S1). Food was removed at egg laying to mimic the natural end of earwigs’ foraging activity [53]. At egg hatching, we discarded all newly emerged larvae from the experiments to trigger a novel ovarian cycle in the mothers and allow their production of a subsequent clutch [31,54]. We then maintained the pairs under the rearing conditions described above until our experiment ended, i.e. either one year after the beginning of our laboratory setup or at the death of the adult males and females. Overall, 3927 of the 4158 (94.4%) tested individuals died within the year following the beginning of our experiments, a value in line with previous data on *F. auricularia* lifespan [51]. Note that recent studies revealed that North American *F. auricularia* encompasses two genetic subspecies with no apparent mixing of their populations [20,35,55]. Although these subspecies were not considered in our analyses (our data were collected before the publication of these genetic analyses), the continuous distribution (unimodal data) of the life history traits measured across populations (Figures 2 to 4) suggests an absence of subspecies-specific values regarding these measurements. The potential co-occurrence of the two subspecies in our data set is thus unlikely to bias our study and its main conclusions.

**Figure 2.**
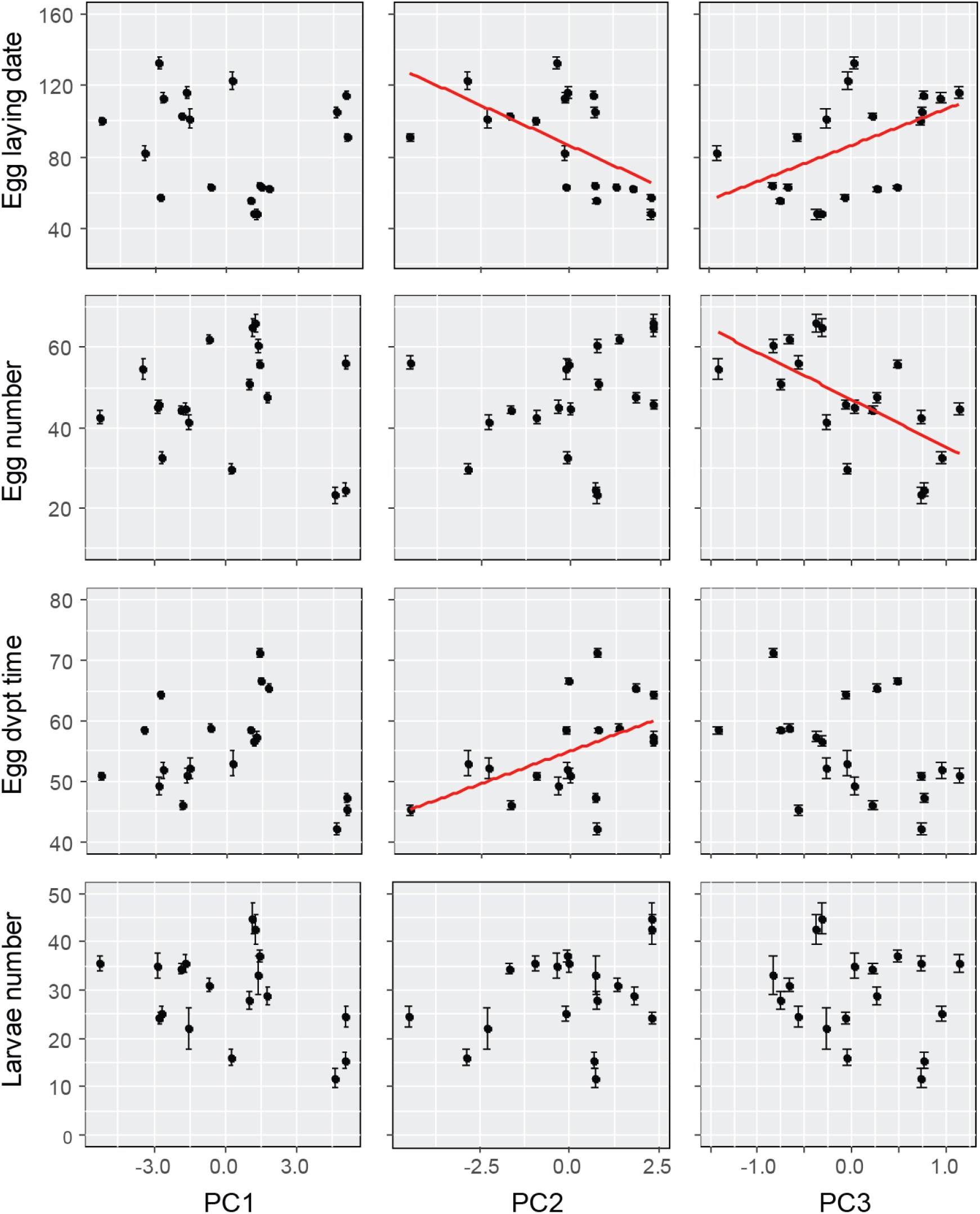
Associations between the thermal regimes (PC1, PC2, PC3) of the 19 populations of origin and 1^st^ clutch parameters. The red lines represent correlations that are significant after FDR correction. Mean values ± SE. Egg laying date was calculated using October ^1st^ as a reference (i.e. as day 0).

### Measurements of the life-history traits

For each mating pair, we measured 13 life-history traits encompassing the properties of the resulting 1^st^ and 2^nd^ clutches (when present), the reproductive strategy and reproductive outcomes of each female, as well as the experimental survival duration of both field-sampled males and females. These properties were obtained by recording the date of egg production, counting the number of eggs produced, calculating the duration of egg development until hatching (in days) and finally counting the number of larvae at egg hatching in both 1^st^ and 2^nd^ clutches (when present). The reproductive strategies and reproductive outcomes of females were obtained by recording whether they were semelparous or iteroparous (i.e. produced one or two clutches in their lifetime, respectively), and by counting the total number of eggs and larvae produced per female during their lifetime. Finally, we measured the experimental survival duration of adults by counting the number of days each male and female survived after October 1^st^ of the year of field sampling.

Although our measurement of survival duration does not necessarily reflect adults longevity, as individuals could have different ages at field sampling (see discussion), it nevertheless provides important insights into the period at which males and females of each population die during the year. Note that 8.1% and 5.4% females from Santa Cruz and Asheville, respectively, produced a third clutch. This third clutch was not considered in the present study, as our experiment ended before their hatching.

### Extraction of mean temperatures and thermal regimes of each population

We extracted the mean monthly temperature of the 19 studied populations using their GPS coordinates (Table 1) and the Worldclim database v2.0 (http://www.worldclim.org/) with a spatial resolution of 30 seconds. The mean temperatures provided by the Worldclim database are calculated over 30 years, from 1970 to 2000. To reduce dimensionality of co-varying temperatures in our data set while characterizing potential thermal regimes of each population without *a priori* definitions of their composition, we then conducted a Principal Component Analysis (PCA) on the set of 12 mean monthly temperatures per population (Table S1). This analysis provided us with 12 orthogonal principal components (PCs), out of which we retained the first three PCs (Table 2; total variance explained = 98.6%). The first component (PC1) was positively loaded by almost all monthly temperatures, therefore positively reflecting the overall mean temperature of a population. The second component (PC2) revealed variation in seasonality between February on one hand, and June, July, and August on the other hand. In particular, high values of PC2 reflected populations with cold February (winter) and warm summer, whereas small values of PC2 reflected populations with warm February (winter) and cold summer. Finally, the third component (PC3) captured variation in seasonality between October and November on one hand, and April and May on the other hand. High values of PC3 therefore characterized populations with cold autumn and warm spring, whereas small values of PC3 characterized populations with warm autumn and cold spring.

**Table 2.** Loadings of the four first principal components (PCs) reflecting combinations of the 12 mean monthly temperatures across populations. The traits having significant loadings on each PC are in bold.

|  | PC1 | PC2 | PC3 | PC4 |
| --- | --- | --- | --- | --- |
| Jan | **0.800** | -0.589 | -0.066 | 0.083 |
| Feb | 0.716 | **-0.668** | 0.131 | 0.139 |
| Mar | **0.844** | -0.486 | 0.216 | 0.048 |
| Apr | **0.949** | -0.140 | **0.267** | -0.082 |
| May | **0.890** | 0.321 | **0.286** | **-0.145** |
| Jun | 0.731 | **0.665** | 0.123 | -0.060 |
| Jul | 0.547 | **0.823** | -0.006 | **0.143** |
| Aug | 0.641 | **0.746** | -0.013 | **0.175** |
| Sep | **0.905** | 0.380 | -0.175 | -0.019 |
| Oct | **0.951** | 0.019 | **-0.292** | -0.064 |
| Nov | **0.931** | -0.174 | **-0.296** | -0.112 |
| Dec | **0.872** | -0.469 | -0.113 | 0.041 |
| Eigenvalues | 8.153 | 3.224 | 0.453 | 0.130 |
| Variance explained (%) | **67.9** | 26.9 | 3.8 | 1.1 |
| Cumulative variance explained (%) | 67.9 | 94.8 | 98.6 | 99.7 |

### Statistical analyses

To test whether *F. auricularia* adapt their life-cycle and life-history traits to North American temperatures, we conducted a series of 12 linear models (LM in R) and one generalized linear model (GLM in R) – see Table 3. In the 12 LMs, the three selected PCs and their interactions were entered as explanatory variables (PC1, PC2 and PC3), whereas the response variable was either egg laying date, egg number, egg development time and larvae number for the 1^st^ or 2^nd^ clutches (for a total of 8 LMs), the total number of eggs or larvae produced, or the survival duration of males or females. Note that both egg laying date and adult survival duration were calculated using October 1^st^ as day 0. In the GLM, the response variable was the ratio of iteroparous females per population, which was entered using the command *cbind* in R (to weight each ratio by the sample size of its population) and fitted to a binomial error distribution corrected for overdispersion. In all our statistical models, the response variables were the mean values of each measured trait per population. They were also checked for homoscedasticity and normality of residuals, as well as simplified stepwise by removing all non-significant interaction terms (all P > 0.05). To correct for inflated Type-I errors due to multiple testing (and provide an experiment-wide Type I error rate of 5%), all *P*-values were adjusted using False Discovery Rate (FDR) correction [56]. All analyses were conducted using the software R v3.5.1 loaded with the packages *raster*, *FactoMineR*, *rsq* and *rcompanion*.

**Table 3.**
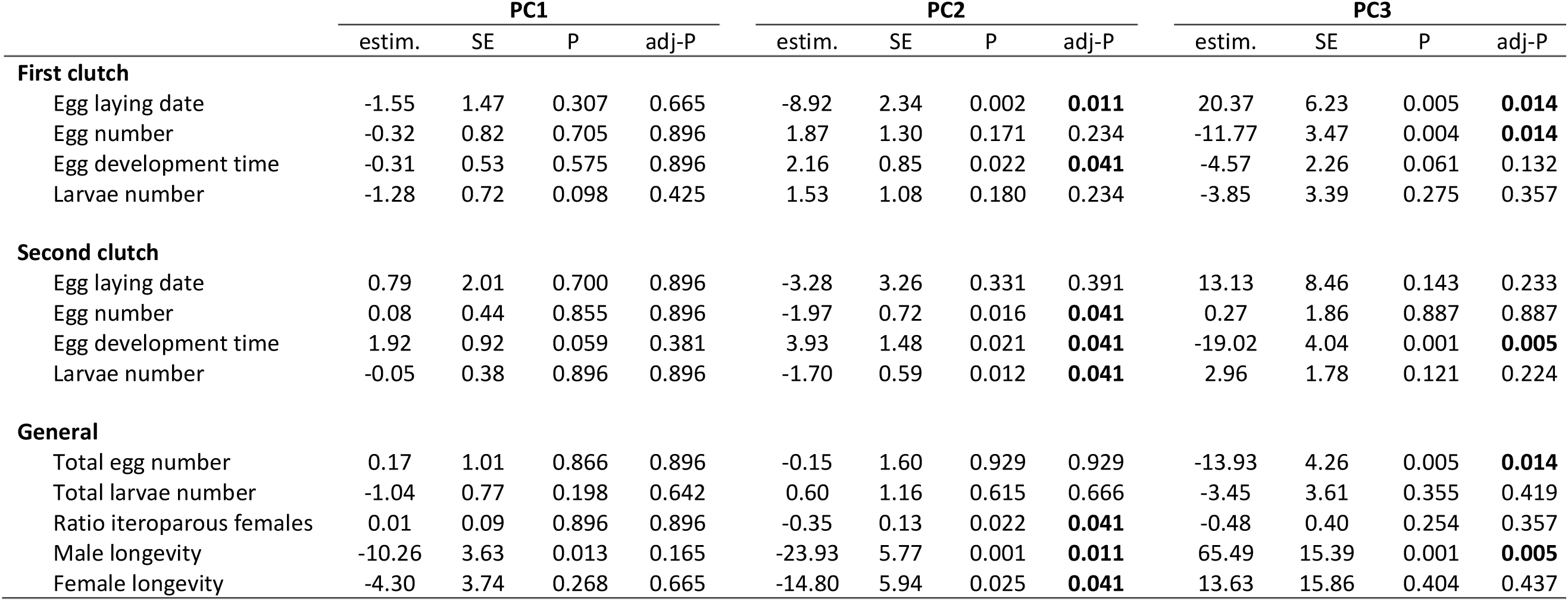
Results of the statistical models on the 13 measured life-history traits. PC1 positively reflects the overall mean temperature of a population. High values of PC2 reflect populations with cold February (winter) and warm summer, and vice-versa. High values of PC3 reflect populations with warm spring and cold autumn, and vice-versa. P-values significant after FDR correction (adj-P) are in bold. Note that FDR correction transforms each P-value in function of its rank of statistical significance in the data set, which can lead to similar corrected p-values. Model estimates (estim) ± standard errors (SE).

## Results

The 19 studied populations greatly varied in their mean temperatures and thermal regimes (Table S1), as well as in the mean values of the 13 traits measured in their sampled individuals (Figures 2 to 4; Tables S2 to S4). Mean monthly temperatures overall ranged from 22.9°C (July in Saluda) to −10.1°C (January in Montreal), while thermal amplitudes over a year ranged from 30.7°C (Montreal) to 7.9°C (Santa Cruz). For the traits measured in the 1^st^ clutches, the mean dates of egg production ranged from 47.8 to 132.6 days after the 1^st^ of October, the mean number of eggs per clutch from 23.2 to 66.0, the mean egg development time from 42.2 to 71.4 days and the mean number of larvae per clutch from 11.6 to 44.8. For the 2^nd^ clutches, the mean dates of egg production ranged from 142.0 to 248.2 days after the 1^st^ of October, the mean number of eggs from 14.0 to 38.4, the mean egg development time from 10.0 to 63.7 days and the mean number of larvae from 0 to 17.7. Finally, the total number of eggs produced ranged from 28.1 to 83.4, the total number of larvae produced from 13.0 to 46.3, the proportion of iteroparous females from 0 to 70.8%, and the survival duration of males and females from 82.0 to 299.8 days and from 146.0 to 322.5 days after the 1^st^ of October, respectively.

Of the 13 measured traits, 10 varied together with the thermal regimes of the population of origin (Table 3). Five of these 10 traits were exclusively associated with PC2 (February-summer temperatures), two traits were exclusively associated with PC3 (autumn-spring temperatures), and three traits were associated with both PC2 and PC3. By contrast, no traits were associated with PC1 (overall mean temperatures). The associations with PC2 revealed that populations with cold February and warm summers (high PC2 values) had females that produced their 1^st^ clutch of eggs earlier and these eggs had longer development time compared to populations exhibiting warm February and cold summers (low PC2 values, Figure 2). Similarly, females from the former populations were less likely to produce a second clutch (i.e. to be iteroparous, Figure 3) and when they did so, their 2^nd^ clutch eggs were less numerous (Figure 3) and showed longer development time (Figure 3). Moreover, females and males from populations with cold February and warm summers lived less long compared to adults from warm February and cold summers (Figure 4). On the other hand, the effects of PC3 reveal that populations exhibiting cold autumns and warm springs (high PC3 values) had females that produced their 1^st^ clutch of eggs later in the season and these eggs were less numerous compared to females from populations with warm autumns and cold springs (low PC3 values, Figure 2). Females from the former populations also had 2^nd^ clutch eggs that exhibited a shorter developmental time (Figure 3), they produced an overall lower number of eggs (Figure 4) and had males with a longer survival duration (Figure 4). By contrast, PC1, PC2 and PC3 did not shape the number of 1^st^ clutch larvae, as well as their total number and the dates of 2^nd^ clutch egg laying (Figures 2, 3 and 4; Table 3).

**Figure 3.**
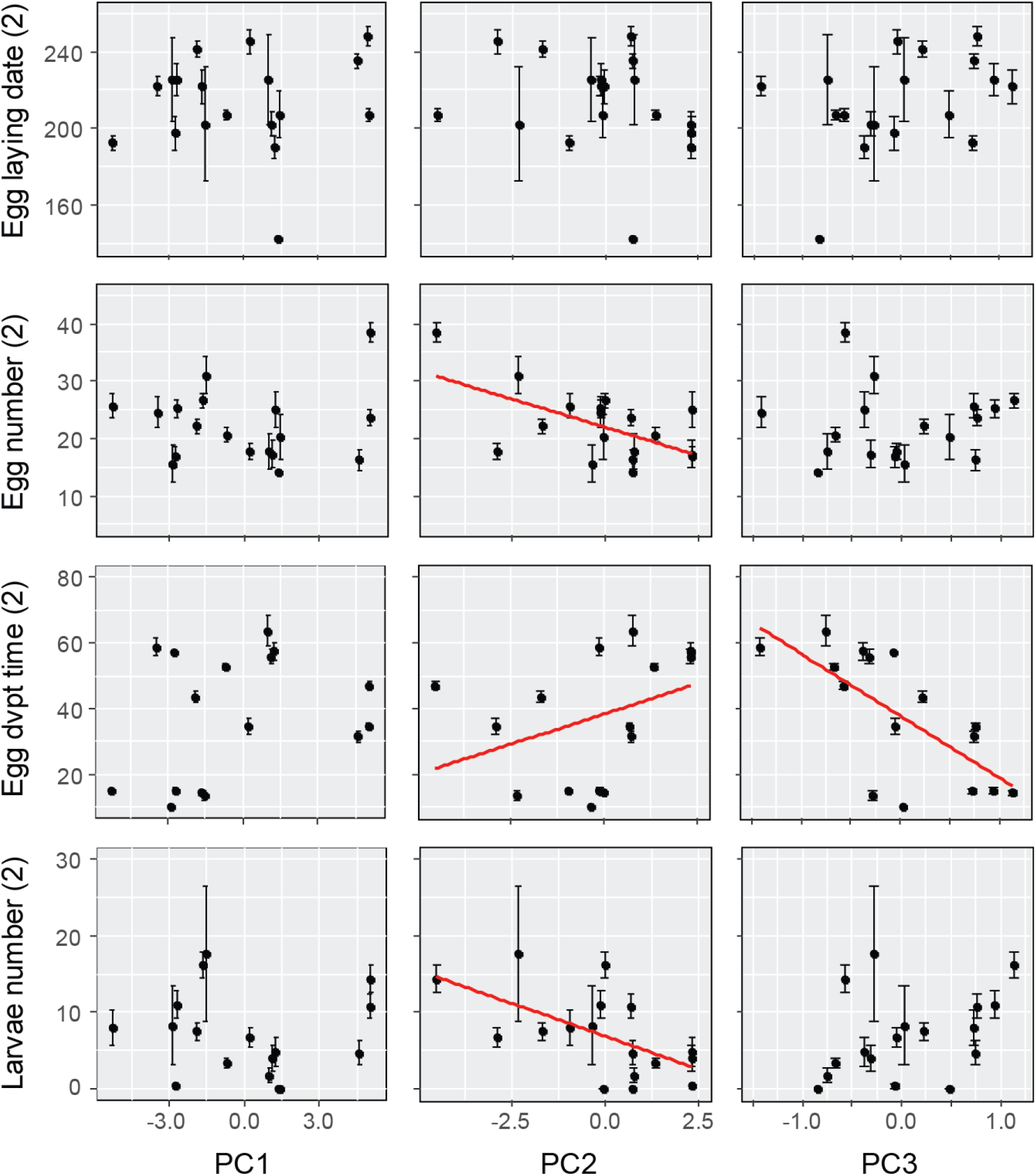
Associations between the thermal regimes (PC1, PC2, PC3) of the 19 populations of origin and 2^nd^ clutch parameters (when produced). The red lines represent correlations that are significant after FDR correction. Mean values ± SE. Egg laying date was calculated using October 1^st^ as a reference (i.e. as day 0).

**Figure 4.**
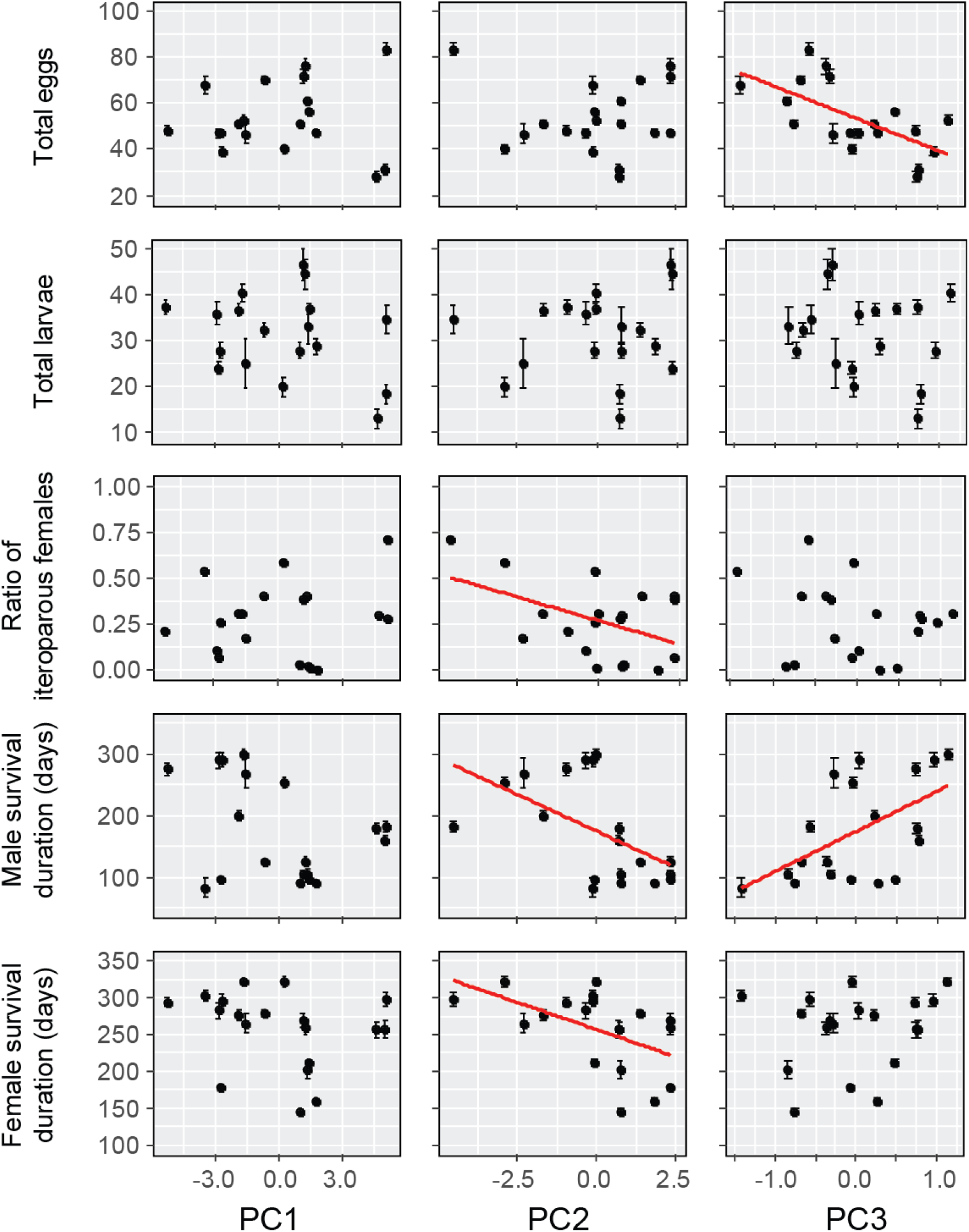
Associations between the thermal regimes (PC1, PC2, PC3) of the 19 populations of origin and females’ reproductive strategies and outcomes, as well as adult’s survival duration. The red lines represent correlations that are significant after FDR correction. Mean values ± SE.

## Discussion

Shedding light on how species successfully adapt to a broad set of environmental constraints is of major importance to improve our general understanding of the mechanisms underlying animal adaptations to climate change. In this study, we demonstrate that the successful invasion of the European earwig across North America came with multiple changes in their life-history traits in response to the thermal regimes (sets of winter-summer and autumn-spring temperatures), but not to the overall mean temperature of the invaded areas. In particular, our data from 19 populations revealed that females changed their timing of first reproduction, their reproductive strategy and investment into egg production when facing different thermal regimes, while experimental survival duration of males and females varied accordingly. By contrast, we found no association between thermal regimes and both the timing of second reproduction and the total number of larvae produced per female.

We first showed that females produced their first clutch of eggs earlier when they came from populations facing warm summers and/or warm autumns (PC2 and PC3, respectively), and were less likely to produce a second clutch in populations with cold February. A plastic response to warm temperatures on egg laying date could be expected in nature: adult earwigs typically develop and mate during summer and autumn, so that warm temperatures during these seasons could accelerate their reproductive physiology (as shown in other insect species, Singh et al., 2018) and thus accelerate egg laying [35]. Similarly, cold Februaries might slow down the development of 1^st^ clutch eggs and thus extend the corresponding period of egg care. This, in turn, might inhibit females’ physiological transformation to produce a second clutch [34,35,43,44]. However, our results were obtained under common garden conditions, which reveals that the observed effects of thermal regime on egg laying dates are not a plastic response to their current environment, but are either due to the environment experienced during their early life development (i.e. before field sampling), or due to an inherited basis that possibly emerged through canalization [12,58]. It has been proposed that traits tightly linked to fitness are more strongly canalized due to past stabilizing selection [59]. Our findings may therefore suggest that the observed changes in the timing of first reproduction and females’ reproductive strategy may have first emerged as a plastic response to the thermal constraints of the different localities, then diverged between populations through canalization and ultimately become inherited traits – all this in a maximum of 100 generations. Further experiments with naïve individuals remain, however, required to rule out an effect of early life experience.

Our data also reveal that thermal regimes are associated with lifetime egg production, but not with lifetime larvae production. In particular, the total number of eggs produced per female decreased with decreasing autumn temperatures, whereas this association vanished with larvae number. This apparent discrepancy suggests that females from populations with the warmest autumns lost a larger number of eggs during egg development. A first explanation could be that these females produced eggs of lower quality and/or were less efficient in egg care, a process that is essential to ensure egg development until hatching in earwigs [40,45]. Whereas both effects should be tested in future studies, previous results may suggest that the second effect is unlikely, as maternal investment in post-hatching care is not population-specific, at least in Europe [36]. Another explanation is that females consumed a larger part of their clutch in populations with the warmest compared to the coldest autumns. Filial egg consumption is a common phenomenon in insects [60] and it has been recently reported in several Dermapteran species, such as the species studied here [41,45] and the maritime earwig *Anisolabis maritima* Bonelli [61]. In the European earwig, this phenomenon has been proposed to reflect an adaptive strategy to limit female weight loss during the period of egg care (i.e. when they stop all other foraging activities) and by doing so, to reallocate resources into post-hatching care and/or into a 2^nd^ oogenesis cycle [35,41]. Given that females lay eggs earlier in populations with the warmest autumns, this increased egg consumption could be an adaptive strategy to limit the cost of tending newly hatched offspring earlier in the season (middle of winter) when food sources are scarce or absent. If this hypothesis holds true, it would suggest that filial egg cannibalism could be a strategy that *F. auricularia* females have evolved to better cope with warmer autumns.

In addition to the above findings, our results show that the survival duration of both males and females was associated with the thermal regime of the population of origin. In particular, females’ and males’ survival duration decreased together with warm summers (and cold Februaries), while male’s survival duration also decreased with warm autumns (and cold springs). The first results may be a by-product of the effect of temperature on their date of egg laying and/or egg hatching. In particular, we showed that females from populations facing warm summers are the first to lay their eggs. Individuals from these populations might thus have been the oldest at the date of our field sampling, therefore leading to the shortest survival duration in our subsequent experiment. Surprisingly, there was a sex-specific effect of spring (and autumn) temperatures on adult survival duration: males lived up to two times longer in populations with warm compared to cold springs (as well as cold compared to warm autumns), whereas this effect was absent in females. This finding may reflect sex-specific sensitivity to high temperatures in terms of, for instance, physiology or expression of costly behaviors. Whereas some physiological traits are known to be sex-specific in this species [52,62], further studies should explore the effects of temperature on the observed differences. Notwithstanding its underlying mechanisms, the long survival duration of males in warm spring populations opens scope for these males to mate with females of the subsequent generation, as well as for a possible involvement of fathers into larva care – a phenomenon reported in other insect species [63]. These two processes remain unknown in the European earwig, but they could be of central importance in their successful adaptation to climate change.

All our results are based on a common garden experiment, a method that is often considered a powerful tool to disentangle the roles of phenotypic plasticity and genetic background on adaptation [15,64,65]. Individuals reared under a common environment are typically expected to exhibit homogenized life-history traits if adaptation is the outcome of phenotypic plasticity, whereas they should exhibit population-specific traits otherwise. Our results are in line with the latter process for the great majority of the measured life-history traits (10 out of 13), therefore suggesting that the observed associations between thermal regimes and life-history traits do not stem from a plastic response to their current environment. Nevertheless, common garden experiments often have some limits: they do not prevent maternal and grand maternal effects, they cannot preclude the possibility of genotype-by-environment interactions on the measured life-history traits, and they are poorly efficient at shedding light on the multiple facets of plasticity (e.g. some traits can be partially plastic, the plastic responses can vary in intensity and slope, and plasticity may become apparent only after certain thresholds) [8,64–66]. These limits can be particularly important here, as maternal effects and harsh environments shape the nature and outcomes of several family interactions in earwigs [42,67–70]. Concluding on the absence or limited role of plasticity in earwigs’ adaptation to North American’ thermal regimes would therefore need further empirical works exploring its multiple facets under several common garden conditions [66], and if present, demonstrating the adaptive value of this apparent plasticity.

To conclude, our results demonstrate that the spread of the European earwigs across North America came with important changes in their life-history traits and life cycle, and that these changes emerged in a maximum of 100 generations. Whereas we show that some of these changes are by-products of novel thermal constraints (timing of first reproduction and female iteroparity), we reveal that others are likely to reflect adaptive strategies to cope with different autumn temperatures (egg production and the possibility of egg cannibalism). Overall, these findings emphasize that adaptation of an insect with a relatively long life-cycle does not necessarily operate in response to the overall mean temperatures of the invaded environments, but to their thermal regimes – i.e. to seasonality and/or mean temperature at a specific time of their life-cycle. Whether the reported adaptations are the product of population-differences in energetic/metabolic constraints experienced by adults during their early development [71,72], and/or the product of an inherited genetic basis that varies with thermal regimes (Levis and Pfennig 2016; Corl et al. 2018; Fox et al. 2019), as well as whether these adaptations are similar across its worldwide distribution [17,20–23] will be investigated in future studies. On a more general level, our findings emphasize that studying invasive species can provide unique data sets to empirically and comprehensively test general predictions on animals’ responses to climate change [4–8,73], and therefore call for their open access to the entire research community - a timely task to which the present study contributes.

## Author contribution statement

JCT designed the experiment, conducted field samplings, and run the experiments. JM analysed the data and wrote the first version of the manuscript. The final manuscript was commented and corrected by all authors.

## Data accessibility

The complete data set and R script are archived in the open data repository Zenodo (https://doi.org/10.5281/zenodo.2652192).

## Acknowledgements

This work is dedicated to the memory of Noelle Tourneur, for her constant support through JCT career. We thank Maximilian Körner, Franck Dedeine and Sylvain Pincebourde, as well as Eric Gangloff, Ben Phillips and Fabien Aubret for their comments on previous versions of this manuscript. We also would like to thank Jean Gingras for his help during laboratory observations and samplings, as well as Michel Vancassel and Maryvonne Forasté for their help in the late 1980s.

